# Bacterial histone HBb from *Bdellovibrio bacteriovorus* compacts DNA by bending

**DOI:** 10.1101/2023.02.26.530074

**Authors:** Yimin Hu, Samuel Schwab, Silvia Deiss, Pedro Escudeiro, Thor van Heesch, Joe D. Joiner, Jocelyne Vreede, Marcus D. Hartmann, Andrei N. Lupas, Birte Hernandez Alvarez, Vikram Alva, Remus T. Dame

## Abstract

Histones are essential for genome compaction and transcription regulation in eukaryotes, where they assemble into octamers to form the nucleosome core. In contrast, archaeal histones assemble into dimers that form hypernucleosomes upon DNA binding. Although histone homologs have been identified in bacteria recently, their DNA-binding characteristics remain largely unexplored. Our study reveals that the bacterial histone HBb (Bd0055) is indispensable for the survival of *Bdellovibrio bacteriovorus*, suggesting critical roles in DNA organization and gene regulation. By determining crystal structures of free and DNA-bound HBb, we unveil its distinctive dimeric assembly, diverging from those of eukaryotic and archaeal histones, while also elucidating how it binds and bends DNA through interaction interfaces reminiscent of eukaryotic and archaeal histones. Building on this, by employing various biophysical and biochemical approaches, we further substantiated the ability of HBb to bind and compact DNA by bending in a sequence-independent manner. Finally, using DNA affinity purification and sequencing, we reveal that HBb binds along the entire genomic DNA of *B. bacteriovorus* without sequence specificity. These distinct DNA-binding properties of bacterial histones, showcasing remarkable similarities yet significant differences from their archaeal and eukaryotic counterparts, highlight the diverse roles histones play in DNA organization across all domains of life.

**Summary:** Histones, traditionally known for organizing and regulating DNA in eukaryotes and archaea, have recently been discovered in bacteria, opening up a new frontier in our understanding of genome organization across the domains of life. Our study investigates the largely unexplored DNA-binding properties of bacterial histones, focusing on HBb in *Bdellovibrio bacteriovorus*. We reveal that HBb is essential for bacterial survival and exhibits DNA-binding properties similar to archaeal and eukaryotic histones. However, unlike eukaryotic and archaeal histones, which wrap DNA, HBb bends DNA without sequence specificity. This work not only broadens our understanding of DNA organization across different life forms but also suggests that bacterial histones may have diverse roles in genome organization.

## Introduction

Histones, a class of highly conserved DNA-binding proteins, are predominantly found in eukaryotes and archaea. These proteins share a characteristic “histone fold”, comprising three α-helices connected by two short strap loops (1,2). In eukaryotes, the four core histones—H2A, H2B, H3, and H4—assemble into an octamer wrapped around by approximately 146 base pairs of double-stranded DNA (dsDNA), forming the nucleosome, the fundamental unit of chromatin (1–3). A unique feature of eukaryotic histones is their unstructured N-terminal tails, which serve as hotspots for post-translational modifications and play pivotal roles in various cellular processes, including transcriptional regulation, DNA repair, replication, and condensation (4,5).

The first archaeal histone homologs, HMfA and HMfB, were identified in the hyperthermophilic archaeon *Methanothermus fervidus* (6). These homologs assemble into dimers that further oligomerize into elongated, nucleosome-like structures, termed hypernucleosomes, upon DNA binding (7–9). Subsequent research has identified one or more histone variants in the majority of archaeal lineages (10–16), including the Asgard superphylum, which comprises the closest prokaryotic relatives of eukaryotes (8,17). Archaeal histones can assemble into both homo- and hetero-oligomers and exhibit varying sizes, with some even possessing N- or C-terminal tails akin to eukaryotic histones (8,18). Additionally, archaeal histone-DNA complexes display a wide variety of histone-oligomerization states. These states are influenced by lateral dimer-dimer interactions and stacking interactions (19), which are not consistently conserved across species. The widespread occurrence and diversity of archaeal histones suggest distinct roles in genome organization and regulation (14,17,20,21).

Notably, histones are also encoded within viral genomes (22,23). Phylogenetically, viral histones are currently positioned between eukaryotic and archaeal histones (24). While many characterized viral nucleosomal histones structurally resemble eukaryotic nucleosome-like structures (25,26), some stack into larger complexes that lack linker DNA, resembling archaeal hypernucleosomes (27).

Historically, it was widely assumed that bacteria lack histone homologs, and any observed instances were thought to have been acquired through horizontal gene transfer (28,29). Instead of histones, bacteria were believed to rely on a multitude of small architectural proteins termed nucleoid-associated proteins (NAPs) (30). These NAPs are sometimes, albeit misleadingly, referred to as “histone-like proteins”. However, NAPs are evolutionarily distinct from histones. Their similarities lie in their abundance, basic nature, and DNA-binding properties. While histones are conserved among eukaryotes, no single NAP is found in all prokaryotes. Despite this broad diversity, NAPs in various prokaryotes perform similar architectural functions by bending DNA, bridging DNA segments, and forming filaments on the DNA (30).

To shed light on the origins of histones and nucleosome-based DNA packaging and regulation, we recently conducted a comprehensive search for bacterial histones and discovered about 600 histone homologs in various bacterial classes, including Actinomycetes, Aquificae, Bdellovibrionia, Cyanophyceae, and Spirochaetia (31). Interestingly, these bacterial histones often co-exist with NAPs, whose structural and functional properties have been extensively studied. NAPs contribute to the local and global structuring of genomes and regulate DNA-associated proteins, including those involved in transcription. Bacterial histones can be categorized into pseudodimeric histones, which consist of two consecutive histone domains on a single chain, and single histone fold proteins with only one domain. Importantly, many bacterial histones contain conserved residues essential for DNA binding in eukaryotic and archaeal histones (Fig. 1A), suggesting that bacterial histones may also bind DNA (31).

**Figure 1.**
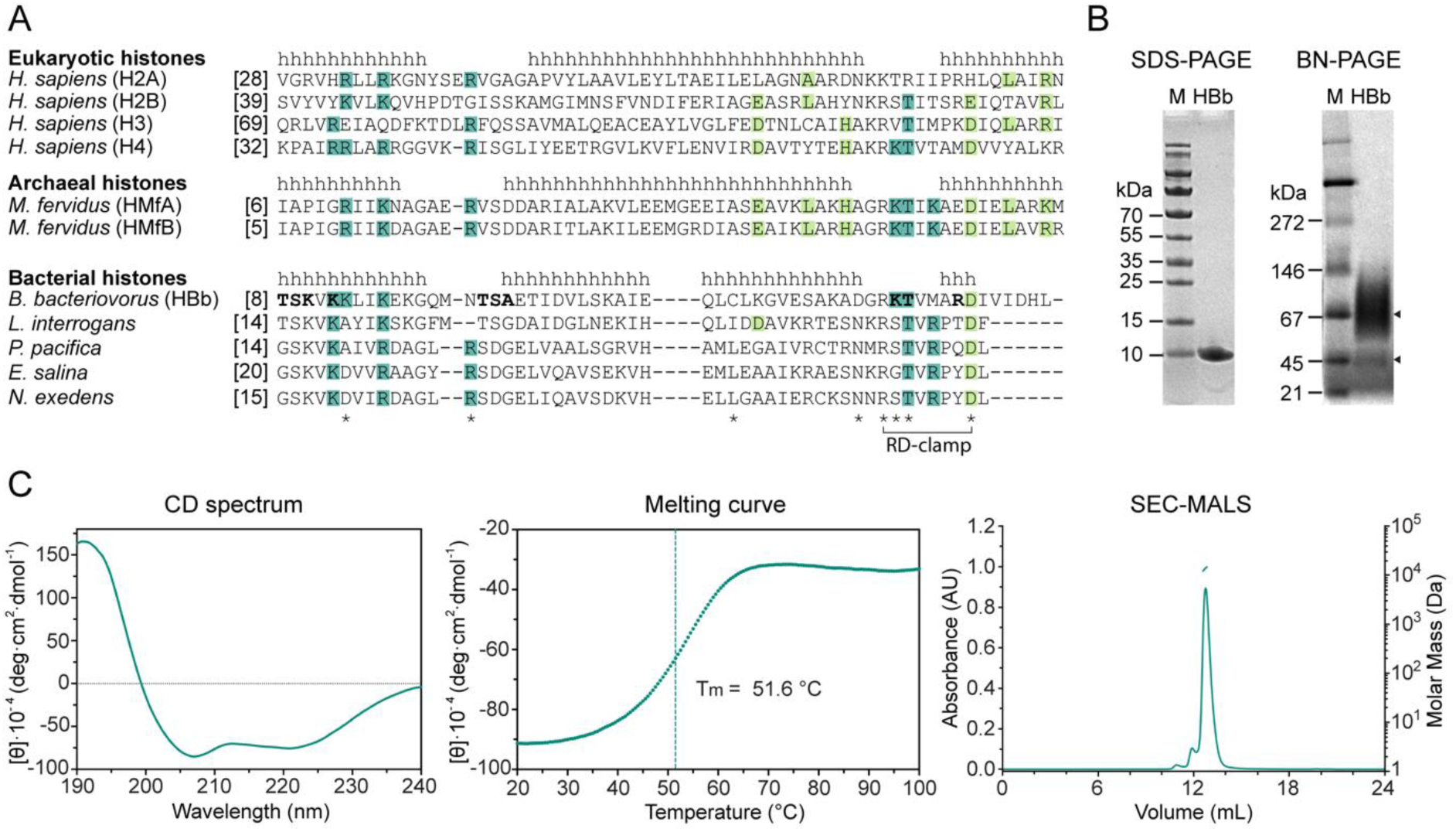
Biophysical characterization of HBb. **A.** Sequence alignment of representative eukaryotic, archaeal, and bacterial histones from *Homo sapiens* (UniProtKB/Swiss-Prot entry for H2A: P04908; H2B: P62807; H3: P68431; H4: P62805), *Methanothermus fervidus* (HMfA: P48781; HMfB: P19267), *Bdellovibrio bacteriovorus* (Q6MRM1), *Leptospira interrogans* (Q8F3E8), *Plesiocystis pacifica* (A6GF99), *Enhygromyxa salina* (NCBI entry WP_052547863.1), and *Nannocystis exedens* (WP_096326703.1). α-Helices are labelled with “h” according to the crystal structures of H2A, HMfB and HBb, respectively. Conserved residues involved in DNA binding and tetramerization are highlighted in dark and light green, respectively. HBb residues involved in DNA backbone interaction according to the solved structures HBb-DNA_1 (PDB: 9EZZ) and HBb-DNA_2 (PDB: 9F0E) are in bold. Residues represented as sticks in Fig. 3 A and B are labelled with **\***. Residues R51 and D58 forming the RD clamp in HBb are connected by a bracket. **B.** SDS-PAGE and BN-PAGE showing purified HBb. Triangles mark HBb on BN-PAGE. **C.** Biophysical characterization of HBb by CD spectroscopy (left: single CD spectrum, middle: thermal melting curve) and SEC MALS (right) for determination of secondary structure, thermal stability and oligomeric state.

In this study, we present crystal structures and biochemical characterization of the free and DNA-bound forms of the histone homolog HBb from the predatory Gram-negative bacterium *Bdellovibrio bacteriovorus*. While the gene encoding HBb is listed in the NCBI databank under the locus tag *BD_RS00255* (formerly *Bd0055*), and even though it was referred to as Bd0055 in a recent publication (32), we propose using the name HBb (<u>H</u>istone of *<u>B</u>*. *<u>b</u>acteriovorus*) to maintain consistency with the standard nomenclature of prokaryotic histones in the literature. Our findings demonstrate that the single histone fold protein HBb forms a homodimer that has the ability to bind and bend DNA. Furthermore, we establish that HBb is essential for maintaining bacterial viability. Our results corroborate many of the observations reported in a recent study by Hocher *et al.* (32), while providing strong evidence for different DNA structuring properties of HBb, suggesting a generic role in global genome organization.

## Materials and methods

Detailed information is provided in the *SI Appendix,* under *Material and Methods*.

### Bacterial strains and cultivation

Genes encoding HBb (*BD_RS00255,* old locus tag: *Bd0055*) and HMfB (GenBank accession number M34778.1) were cloned for recombinant protein expression in *E. coli*. *B. bacteriovorus* HD100 (DSM 50701, DSMZ) was cultured to construct a chromosomal *BD_RS00255* deletion strain using *E. coli* Top10 as prey cells.

### Cloning, plasmids and synthetic DNA

Plasmids pETHis1a (33) and pET28a were used as expression vectors for HBb and HMfb protein production, respectively. For chromosomal deletion of *BD_RS00255* in *B. bacteriovorus*, the suicide vector pT18mobSacB (Addgene plasmid # 72648; http://n2t.net/addgene:72648; RRID:Addgene_72648) was used. Synthetic oligonucleotides (Merck) used in this study are listed in Table S1.

### Deletion of *BD_RS00255*

Attempts to delete *BD_RS00255* in *B. bacteriovorus* HD100 were conducted as previously described, with slight modifications (34,35).

### Protein expression and purification

HBb, HMfB, and FLAG-HBb were recombinantly expressed in *E. coli* Mutant56(DE3) (36) and purified to homogeneity using affinity and size-exclusion chromatography.

### Circular dichroism (CD) spectroscopy

CD spectroscopy was performed to analyze the secondary structure and thermal stability of purified HBb protein using a Jasco J-810 spectrometer (JASCO).

### Size exclusion chromatography coupled with multi-angle light scattering (SEC-MALS)

SEC-MALS was performed to measure the mass of purified HBb in solution, using a 1260 Infinity II HPLC system (Agilent) coupled to a miniDAWN TREOS and Optilab T-rEX refractive index detector (Wyatt Technology).

### Crystallization, data collection, and structure determination

Crystallization trials of HBb were performed in presence and absence of DNA. Crystals were obtained and structures solved for the free and two different DNA-bound forms. Detailed methods are described in detail in *SI Appendix.* Data processing and refinement statistics are listed in Table S3, and the coordinates and structure factors have been deposited in the PDB under accession codes 8CMP (free HBb), 9EZZ (HBb-DNA_1) and 9F0E (HBb-DNA_2).

### Microscale thermophoresis (MST)

The DNA binding affinities of HBb to the 80 bp (9,37) and 80 bp-GC50 oligonucleotides were measured using the Monolith NT.115 instrument (NanoTemper Technologies). MST data were analyzed using the associated MO.Control software.

### DNA binding analysis

Binding of HBb and HMfb to DNA was comparatively analyzed using various assays, including electrophoretic mobility shift assay (EMSA), Micrococcal Nuclease (MNase) digestion assay, DNA topology assay, Ligase-mediated circularization assay, and tethered particle motion (TPM) experiments.

### DNA affinity purification sequencing (DAP-seq)

Preparation of fragmented DNA libraries from genomic DNA of *B. bacteriovorus* HD100, subsequent DNA affinity purification, and next-generation sequencing were conducted as previously described, with some modifications (38). Data were processed with the nf-core/chipseq (version 2.0.0) pipeline (39), under Nextflow (version 23.04.2).

### Modeling of the HBb-DNA complex

Molecular dynamics (MD) simulations of binding of the HBb dimer to DNA were performed using a structural model of an HBb dimer generated based on the two crystal structures of HBb with bound DNA (PDB: 9EZZ and 9F0E) and an AlphaFold2 (AF2) prediction (40). Methods and software used for MD simulations are described in detail in the *SI Appendix, Material and Methods*.

## Results

The majority of bacterial histone homologs that we have identified in a recent bioinformatics analysis originate either from sparsely characterized bacteria of marine origin or from environmental samples (31). The predatory bacterium *B. bacteriovorus* represents one of the few organisms identified in the analysis that is both culturable in the lab and amenable to genetic manipulation. For these reasons, we selected the histone protein HBb of *B. bacteriovorus* (UniProt entry Q6MRM1) for comprehensive functional and structural characterization.

### The histone HBb from B. bacteriovorus binds DNA in vitro

We overexpressed HBb recombinantly in *Escherichia coli* and purified the protein to homogeneity. Its purity was confirmed through SDS-PAGE, which displayed a single band corresponding to a molecular weight of 10 kDa, in line with the theoretical molecular weight of 7 kDa (Fig. 1B). BN-PAGE separation yielded two bands (Fig. 1B), suggesting the presence of diverse oligomers. Nonetheless, SEC-MALS analysis showed only a major peak with a molecular weight of 14 ± 0.4 kDa, indicating a dimer (Fig. 1C). Circular dichroism (CD) spectroscopy indicated HBb to be an α-helical protein with flexible regions that cooperatively unfolds when heated, exhibiting a melting temperature of 51.6°C (Fig. 1C).

We investigated the potential interaction of HBb with DNA using an electrophoretic mobility shift assay (EMSA) with an 80 bp DNA fragment that has a high affinity for HMfB (9,37). We observed a band shift, indicative of HBb-DNA complex formation. This was markedly different from the well-defined protein-DNA bands formed by hypernucleosomal histones, such as HMfB (Fig. 2A). The HBb-DNA complexes appeared as smeared bands, suggesting varied conformations, stoichiometries, or potential protein dissociation. We quantified the binding affinity of HBb to the 80 bp DNA fragment and a DNA fragment of random sequence with a GC content of 50% (80 bp-GC50) using microscale thermophoresis (MST) (Fig. 2B and Table S2). Our results showed that HBb binds nonspecifically to DNA, with an affinity in the lower micromolar range, comparable to HMfB (41–43).

**Figure 2.**
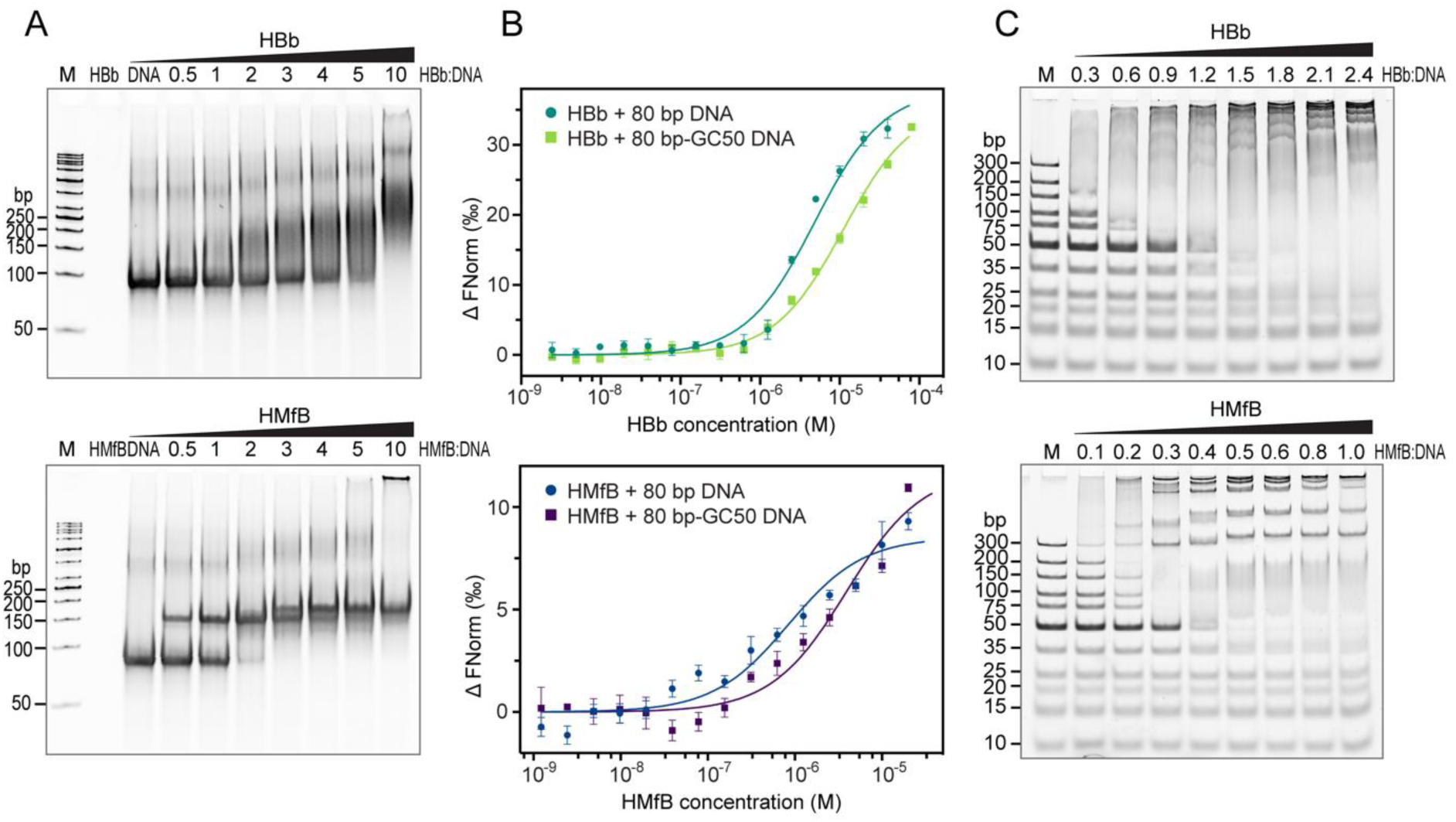
HBb binds DNA. **A.** EMSA with the 80 bp DNA fragment in the presence of HBb and HMfB as a control. The molar ratios of protein:DNA in lanes 4-10 are indicated. **B.** MST curves showing binding of HBb to the 80 bp DNA fragment and the 80 bp-GC50 DNA fragment (10 nM), as compared to HMfB. **C.** EMSA of HBb and HMfB as a control with the GeneRuler Ultra Low Range DNA Ladder (ThermoFisher Scientific). For lanes 2-9, the protein/DNA mass ratios are labelled.

To estimate the size of the DNA binding site of HBb, we performed another EMSA using a DNA ladder (Fig. 2C). Stable HBb-DNA complexes were observed with fragments equal to or larger than 35 bp. Although no band shift was observed for the 20 bp and 25 bp fragments, these fragments appear blurred, suggesting that HBb forms unstable complexes with these smaller fragments. In this EMSA ladder assay, HMfB binds to fragments larger than 35 bp, in agreement with previous EMSAs performed on small substrates (44).

### HBb crystal structure

Crystallization trials were conducted with purified HBb, and within a few days, we obtained crystals that diffracted to atomic resolution. The best dataset could be processed to a resolution of 1.06 Å and the structure was solved by molecular replacement using the archaeal histone HMk (PDB: 1F1E) as a search model. Except for the first three and last three amino acids, the full chain is clearly resolved in the electron density map. The asymmetric unit contains a single chain of HBb, forming a dimer through crystallographic symmetry. HBb exhibits a characteristic dimeric histone fold, with monomers consisting of three α-helices (α1, α2, α3) connected by two loops (l1 and l2) (Fig. 3A). In each monomer, the longer central helix α2 is flanked at both ends by the shorter helices α1 and α3. Dimerization occurs through the antiparallel alignment of the central α2-helices of the two monomers at an angle of approximately 25°. The dimer is stabilized primarily by hydrophobic contacts along the interface, involving residues L31, I35, and L38 in α2, as well as contributions from α1 (e.g. V11) and α3 (e.g. I59).

**Figure 3.**
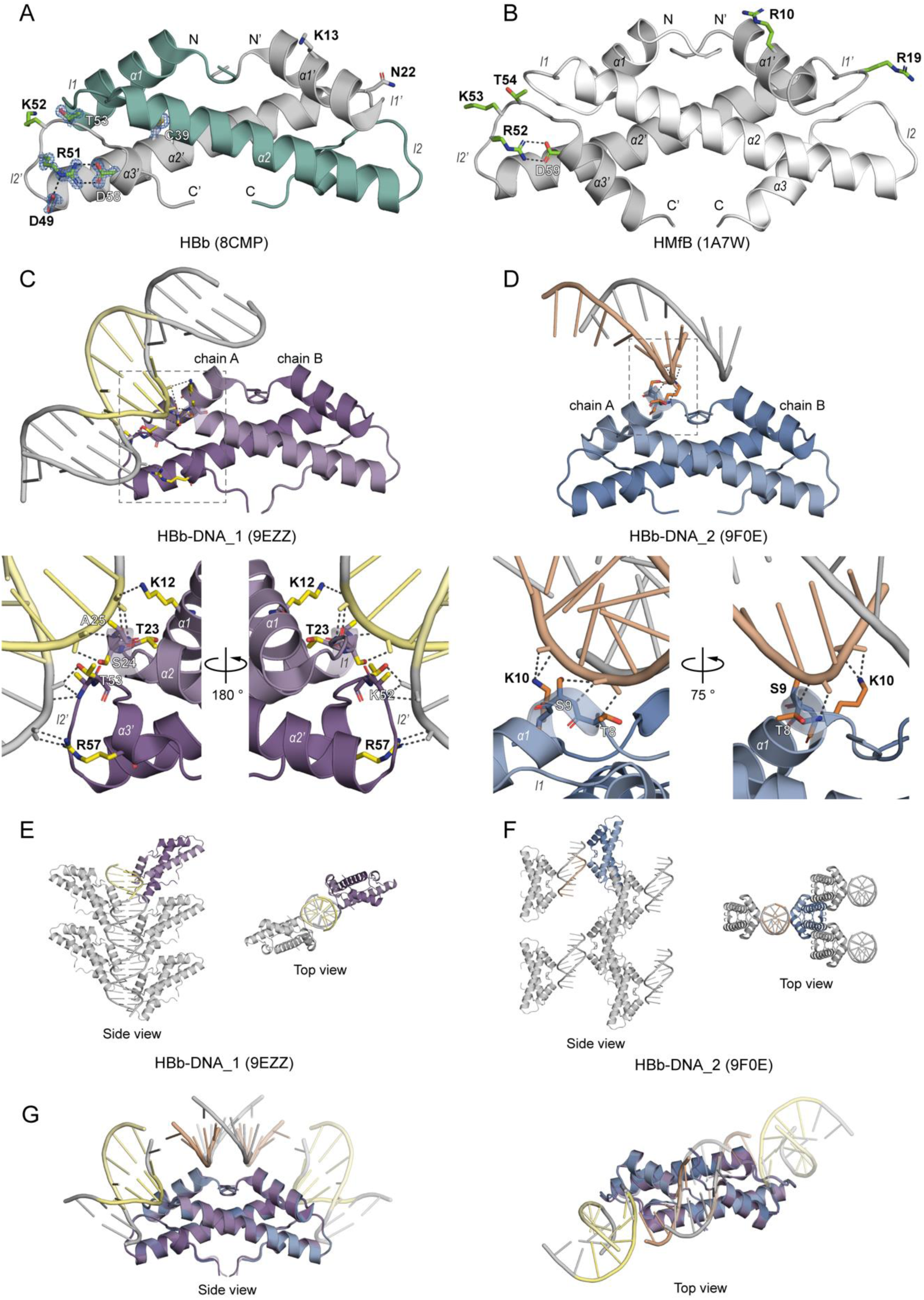
Crystal structures of HBb in its free and DNA-bound forms. **A.** Crystal structure of the HBb dimer (PDB: 8CMP) in cartoon representation. Selected residues are shown as sticks and conserved residues predicted to be involved in DNA binding or stabilization of the histone fold are in green. The salt bridges formed between D49, R51, and D58 are indicated as dashed lines. A representative 2Fo – Fc electron density map is shown for selected residues at a contour level of at 2.0 σ. **B.** Crystal structure of the HMfB dimer (PDB: 1A7W) in cartoon representation for comparison. Conserved residues playing a role in DNA binding or stabilization of the histone fold are depicted as sticks. **C.** Crystal structure of HBb-DNA_1 (PDB: 9EZZ) in cartoon representation with residues involved in DNA binding shown as sticks. The dashed frame marks the enlarged image section. **D.** Crystal structure of HBb-DNA_2 (PDB: 9F0E) in cartoon representation with DNA binding residues shown as sticks. The enlarged image section is marked by a dashed square. **E.** Crystal packing of HBb-DNA_1 with selected symmetry mates generated within 4 Å. **F.** Crystal packing of HBb-DNA_2 with selected symmetry mates generated within 20 Å. **G.** Superposition of structures HBb-DNA_1, HBb-DNA_2, and their symmetrical mates, with DNA fragments trimmed for visualization. In all panels, the contents of a single asymmetric unit are shown in color and selected symmetry mates in gray. Hydrogen bonds and salt bridges are indicated as dashed lines.

In contrast to the archaeal histone HMfB (Fig. 3B), the α2-helix of HBb is one turn shorter, and α3 comprises only a single helical turn. Like HMfB, HBb features a conserved intramolecular salt bridge, known as the RD clamp, involving residues R51 and D58 (Fig. 1A, 3A, B). In HBb, this motif is augmented by D49, which forms an additional salt bridge with R51, potentially stabilizing the conserved KT motif in loop l2. This KT motif in HMfB is known to be involved in DNA interactions. Four of the six DNA-binding residues in HMfB, including K52 and T53, are conserved in HBb (Fig. 1A), indicating a potential for DNA binding.

### Crystal structure of HBb bound to DNA

Similar to its free form, HBb successfully crystallized in complex with a randomized 20 bp DNA fragment (20 bp-GC50) within a few days, yielding the two distinct crystal forms HBb-DNA_1 and HBb-DNA_2. These could be processed to a resolution of 1.95 Å (HBb-DNA_1) and 1.85 Å (HBb-DNA_2), and solved by molecular replacement using the free form of HBb as a search model. Both structures contain one HBb dimer in the asymmetric unit, with all residues apart from the termini clearly resolved in the electron density. Additionally, clear electron density for short DNA fragments were apparent in both structures, albeit in different positions.

For HBb-DNA_1, the asymmetric unit contains a 5 bp dsDNA fragment, which forms an infinite dsDNA helix throughout the crystal via crystallographic symmetry (Fig. 3C). Both monomers of the HBb dimer interact with the minor groove of this DNA (Fig. 3C). Specifically, residues K12 in helix α1, T23 and S24 in loop l1, and A25 in helix α2 of one monomer, as well as K52 and T53 in loop l2 and R57 in helix α3 of the other monomer, form an interaction network with the phosphate groups of the DNA backbone through their side chains and backbones. This structure is largely consistent with the structure recently published by Hocher et al. (PDB: 8FW7), showing the same mode of HBb binding to DNA (Fig. S1) (32). However, the two structures differ in their crystal packings, notably in the number and arrangement of HBb dimers along the DNA (Fig. 3E, Fig. S1).

In contrast, HBb-DNA_2 reveals a complementary DNA binding interface. Here, the asymmetric unit contains a 9 nucleotide (nt) long single-stranded DNA fragment, which forms dsDNA fragments via crystallographic symmetry (Fig. 3D). These fragments are bound by HBb via an additional interface of the minor groove. This involves interactions between residues T8, S9, and K10, all located in the first turn of the helix α1 and conserved in other bacterial histone homologs, and the DNA phosphate backbone (Fig. 3D). The crystal packing in HBb-DNA_2 markedly differs from that in HBb-DNA_1 in terms of the arrangement of the HBb dimers and DNA, containing only short DNA segments (Fig. 3E, F).

The superposition of the HBb-DNA_1 and HBb-DNA_2 structures illustrates the location of the combined HBb DNA binding interfaces (Fig. 3G), resembling histone-DNA complexes found in eukaryotes and archaea. The engagement of these interfaces suggests that the DNA undergoes bending upon binding with the HBb dimers.

### HBb bends DNA *in vitro*

To further substantiate the potential bending of DNA upon HBb binding, as suggested by our crystal structures, we conducted tethered particle motion (TPM) experiments with a linear 685 bp DNA fragment to determine the extent to which HBb alters DNA conformation upon binding. TPM is a single-molecule technique that allows for real-time observation of DNA-protein interactions. In this method, a DNA molecule is tethered at its ends on a bead and a glass surface, respectively, and the motion of the bead serves as a proxy for the conformational state of the DNA. TPM detects conformational changes of the DNA molecule as a function of the protein concentration, measured by the change in root mean square (RMS) displacement of the bound bead. Proteins that compact the DNA reduce the distance between the ends of the DNA, which is measured as a decrease in RMS. In the presence of HBb, the RMS values decrease significantly and reach saturation at ≈110 nm (Fig. 4A). This value is comparable to the TPM values we measured previously for other DNA-bending proteins, such as *E. coli* HU and *Sulfolobus solfataricus* Sul7 and Cren7 (45,46). However, it is higher than the values reported for the archaeal hypernucleosome-forming histones HMfA and HMfB (19), strongly suggesting that HBb bends DNA rather than wrapping it.

**Figure 4.**
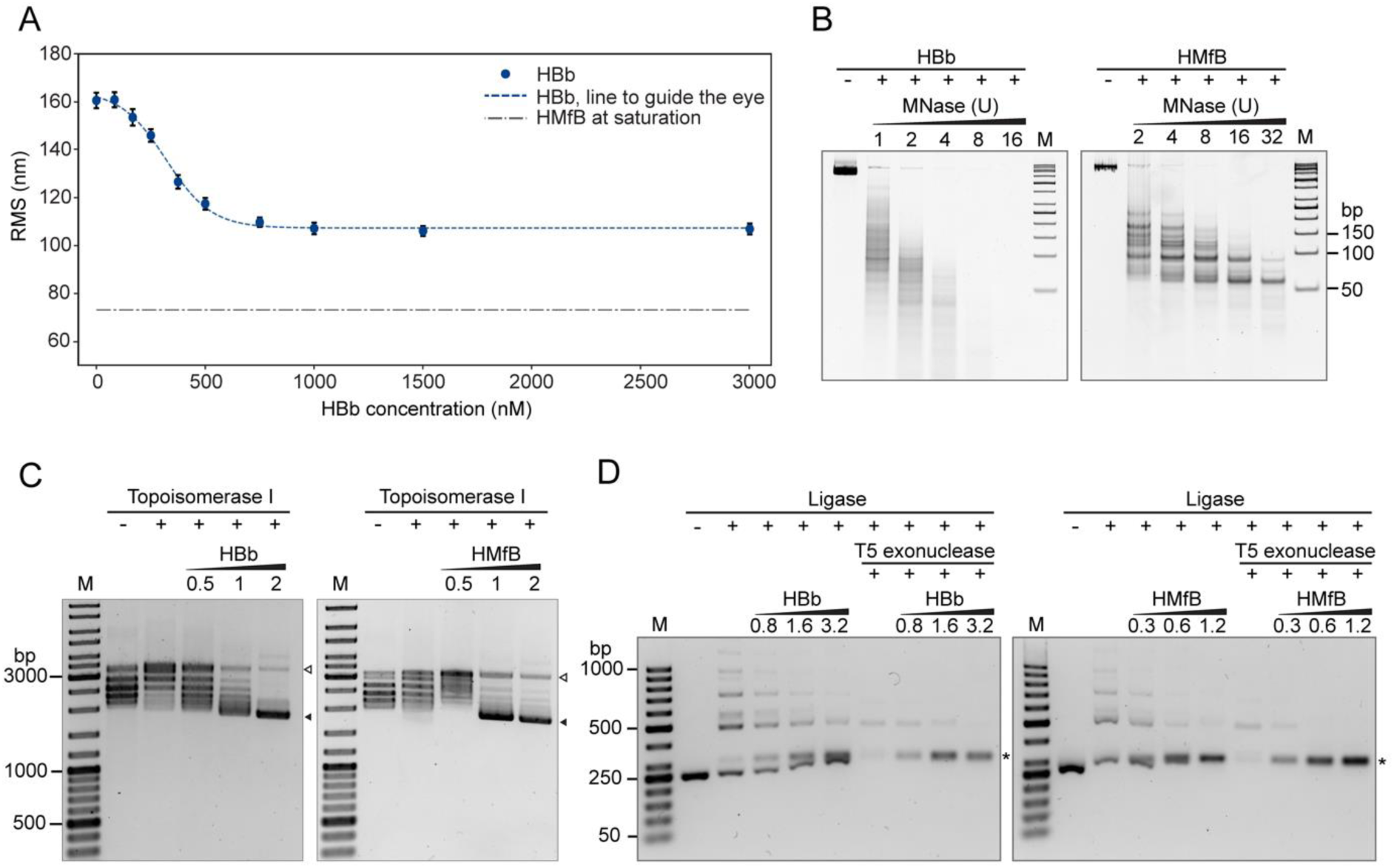
HBb is a DNA bender. **A.** TPM experiment of HBb with 685 bp DNA. All measurement points are averages of triplicate measurements, with error bars indicating standard deviations. As reference, the RMS value obtained for hypernucleosome structure formed with HMfB is shown as the gray dash-dotted line (19). **B.** MNase digestion of 685 bp DNA following incubation with HBb and HMfB. Increasing amounts of MNase in the assay are indicated in Units. **C.** DNA topology assay with relaxed pUC19 in the presence of HBb in comparison to HMfB. Protein:DNA mass ratios were labeled in the figure. The bands showing relaxed (△) and supercoiled pUC19 (▴) are marked. **D.** Ligase-mediated circularization assay of a 240 bp DNA in the presence of HBb and HMfB. Samples are shown before and after T5 exonuclease digestion. Protein:DNA mass ratios are labeled. Monomeric DNA rings are indicated (*).

We further characterized the DNA binding properties of HBb by performing well-established assays, including micrococcal nuclease (MNase) digestion, DNA topology, and ligase-mediated circularization assays (44,47–49). In the MNase digestion assay, MNase is used to digest DNA that is not protected by bound proteins. Unlike the positive control HMfB, which tightly packs DNA and protects multiples of 30 bp fragments from endo-exonuclease digestion (11), the binding of HBb to DNA, similar to NAPs (50,51), offered no such protective effect (Fig. 4B). In the DNA topology assay, we analyzed whether the binding of HBb alters the geometry of DNA. Such changes affect the supercoiling state of DNA in the presence of Topoisomerase I and can be visualized by separation of the DNA in an agarose gel. Here, both HBb and HMfB constrain supercoils in a concentration-dependent manner (Fig. 4C). Using a ligase-mediated circularization assay, we next examined whether this change in DNA geometry results from DNA bending. DNA-bending proteins effectively render small DNA fragments more flexible, thus promoting the formation of small circular DNA during a ligation reaction. The unambiguous identification of circular DNA is possible by treatment of the ligation product with T5 exonuclease, followed by separation through gel electrophoresis. In the assay, fewer linear fragments and more 240 bp circles formed upon addition of increasing amounts of HBb to 240 bp linear DNA and T4 DNA ligase, indicative of DNA bending induced by HBb (Fig. 4D).

### MD simulations support the DNA bending model

To further validate our hypothesis that HBb functions as a DNA-bending histone dimer, we turned to Molecular Dynamics (MD) simulations. For the starting structure, we combined the HBb-DNA_1 and HBb-DNA_2 crystal structures to create a complete DNA molecule bent around the HBb dimer (Fig. 5A, S2). With this starting structure, we performed two 500 ns MD simulations (Supplementary videos 1 and 2). During both 500 ns simulations, the DNA remains bent by HBb with a bending angle around 84 ° ± 3.4 ° across 36 base pairs (Fig. 5B). Due to the bending, HBb compacts DNA as illustrated by the lower end-to-end distance compared to the contour length of the DNA (Fig. S3A). HBb interacts exclusively with the backbone of the DNA, suggesting nonspecific DNA binding by HBb. Important DNA-binding residues are N22, R57, S9, T53, T8, S24, K10, K12, K52, and K16, ordered from the highest number of DNA contacts (14) to the lowest (5) (Fig. 5C, S3B, S3C, S3D). Note that residues with less than 5 contacts are not considered important. These residues interact exclusively at the minor grooves of the DNA and are conserved in related bacterial histones from Bdellovibrionota, Elusimicrobiota, Spirochaetota, Planctomycetota, Myxococcota, and Chlamydiota (Fig. 1A) (52). Bending by HBb causes severe deformations in the minor and major grooves (Fig. S3E, S3F). The minor grooves where HBb binds with its l1-l2 loop and α1 helix are 0.3 nm narrower, while the major grooves are 0.6 nm wider. In contrast, the major groove between the l1-l2 loop and α1 helix binding sites narrows by 0.5 nm, while the minor groove widens by 0.2 nm. This periodic variation between wide and narrow major and minor groove widths is similar to what is observed in canonical nucleosome histones (53). Overall, the DNA-bending structure of the HBb dimer is highly reminiscent of both the archaeal and eukaryotic histone dimers (Fig. 5D, 5E). Both histones contact the DNA at their l1-l2 loops and α1 helix and interact primarily with the DNA backbone. The bending angle for both HBb and canonical histone HMfB is 84 ° across 36 base pairs (9). However, as HBb does not form larger wrapping oligomers like canonical histones, it will provide less DNA compaction than canonical histones, as illustrated by our TPM results (see “HBb bends DNA *in vitro*”).

**Figure 5.**
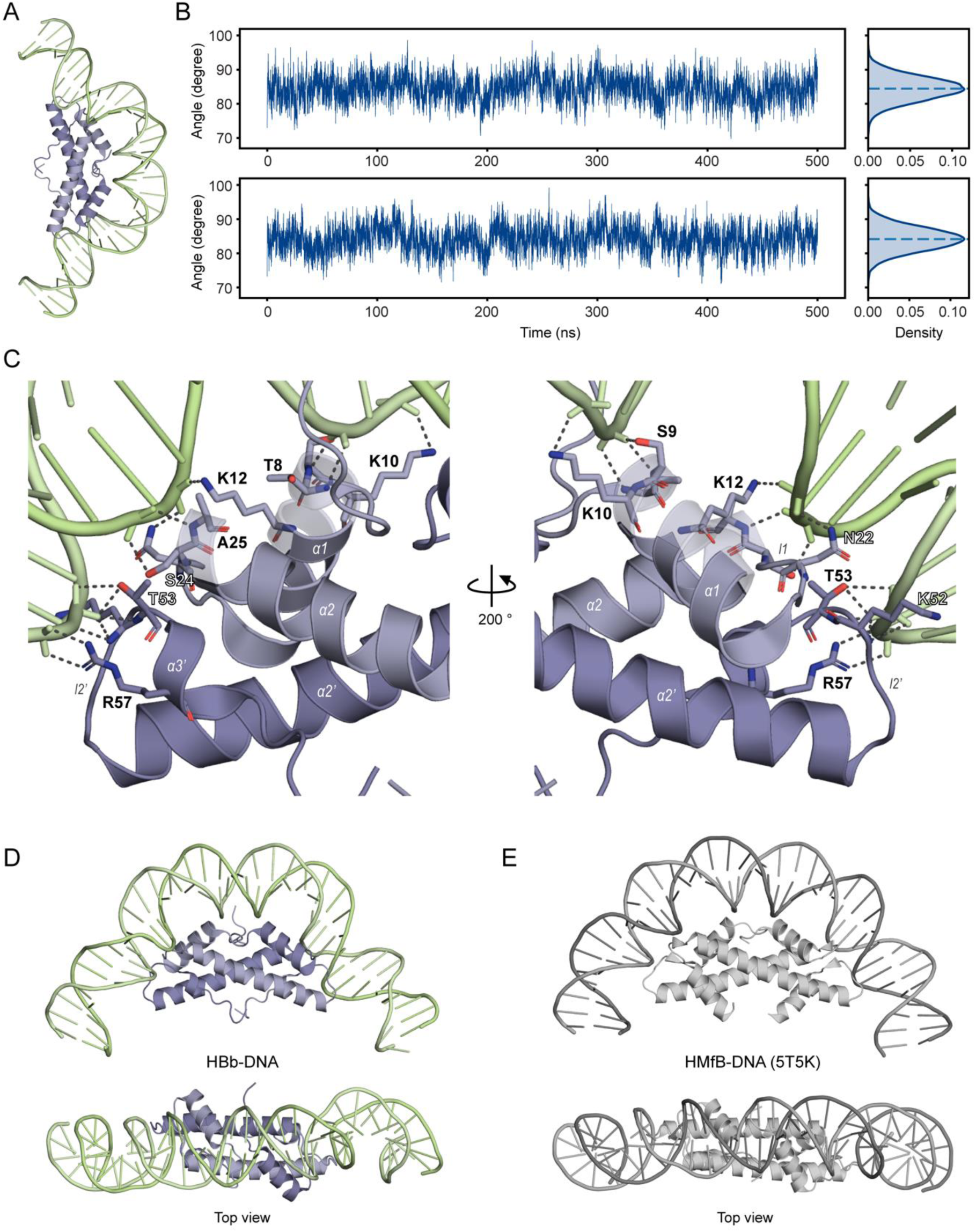
MD simulations supporting DNA bending by HBb. **A.** The HBb-DNA starting model used for MD simulation. The HBb-DNA model was constructed based on HBb-DNA crystal structures followed by energy minimization, and temperature and pressure equilibration. **B.** Plots showing the bending angle of the DNA during two independent 500 ns simulations. **C.** The enlarged image section of the DNA-bound HBb structure obtained from MD simulations. The structure shown is the central structure of the most common structure cluster in the last 100 ns of the simulations. Residues with a large number of contacts to DNA are shown as sticks. **D.** The side view and top view of the cartoon representation of the central structure of the most common cluster of the simulations. E. The side view and top view of a section of HMfB bound to DNA (PDB: 5T5K). The cartoon representation shows an HMfB dimer with a 38-bp dsDNA wrapping around it.

### Nonspecific binding of HBb to the *B. bacteriovorus* chromosome is essential for viability

The bacterium *B. bacteriovorus* is a well-studied model organism with a predatory life cycle. To further understand the functional significance of HBb, we aimed to replace the gene encoding HBb, *BD_RS00255*, with a kanamycin resistance cassette using the suicide vector pT18mobSacB. We achieved successful integration of the plasmid into the chromosome but failed to select for a second crossover event. This suggests that HBb is essential for *B. bacteriovorus* growth and corroborates recent findings (32).

In a subsequent step, we investigated whether HBb has DNA sequence specificity and possibly functions as a transcription factor. To this end, we conducted DAP-seq on a chromosomal scale, a recently developed method to identify genome-wide binding regions of DNA-binding proteins (38). We coupled purified FLAG-HBb to magnetic beads coated with FLAG antibodies and incubated them with a genomic *B. bacteriovorus* DNA library. Protein-bound DNA fragments were isolated, purified, amplified, and subjected to next-generation sequencing. We deviated from the original protocol by expressing FLAG-HBb in *E. coli* rather than in its natural host *B. bacteriovorus*, as this did not yield a sufficient amount of protein. Moreover, we incubated the magnetic beads directly with purified protein instead of cell extract to minimize the risk of nonspecific binding of *E. coli* proteins. DAP-seq analysis was performed for two independently prepared genomic DNA libraries with a fragment size of approximately 200 bp, representing two biological replicates. Samples containing only DNA incubated with anti-FLAG beads in the absence of protein served as controls. For these samples, as determined by Bioanalyzer analysis, the quantity of recovered amplified DNA was less than the amount of primer dimers formed during the amplification. Since this confirms the absence of non-specifically bound DNA, sequencing of these samples was omitted.

Plotting the raw data from DAP-seq analysis on the chromosome map of *B. bacteriovorus* shows HBb binding to both genic and intergenic regions randomly throughout the genome (Fig. 6). After peak-calling analysis (as described in *Supplemental Materials and Methods*), we identified a total of 475 statistically significant HBb-binding sites for DAP-seq sample 1 and 455 for DAP-seq sample 2, respectively, distributed across the chromosome (Fig. 6A). Most of these peaks are between ∼230 bp and ∼660 bp in length reaching maximum lengths of 2,416 bp and 1,747 bp for DAP-seq sample 1 and 2, respectively (Fig. S4). This indicates that while HBb binds to DNA throughout the entire genome, certain regions are notably enriched with HBb dimers, which are likely spaced along the DNA. Analysis of the peaks in relation to the GC content of the DNA fragments covered yielded a value of approximately 50%, which corresponds to the GC content of 50.5% of the *B. bacteriovorus* HD100 genome (54) (Fig. S5). This suggests that HBb binds DNA without preferring GC-rich or AT-rich regions. The search for special sequence motifs proved to be infeasible given the broad length distribution and random pattern of binding sites across the chromosome.

**Figure 6.**
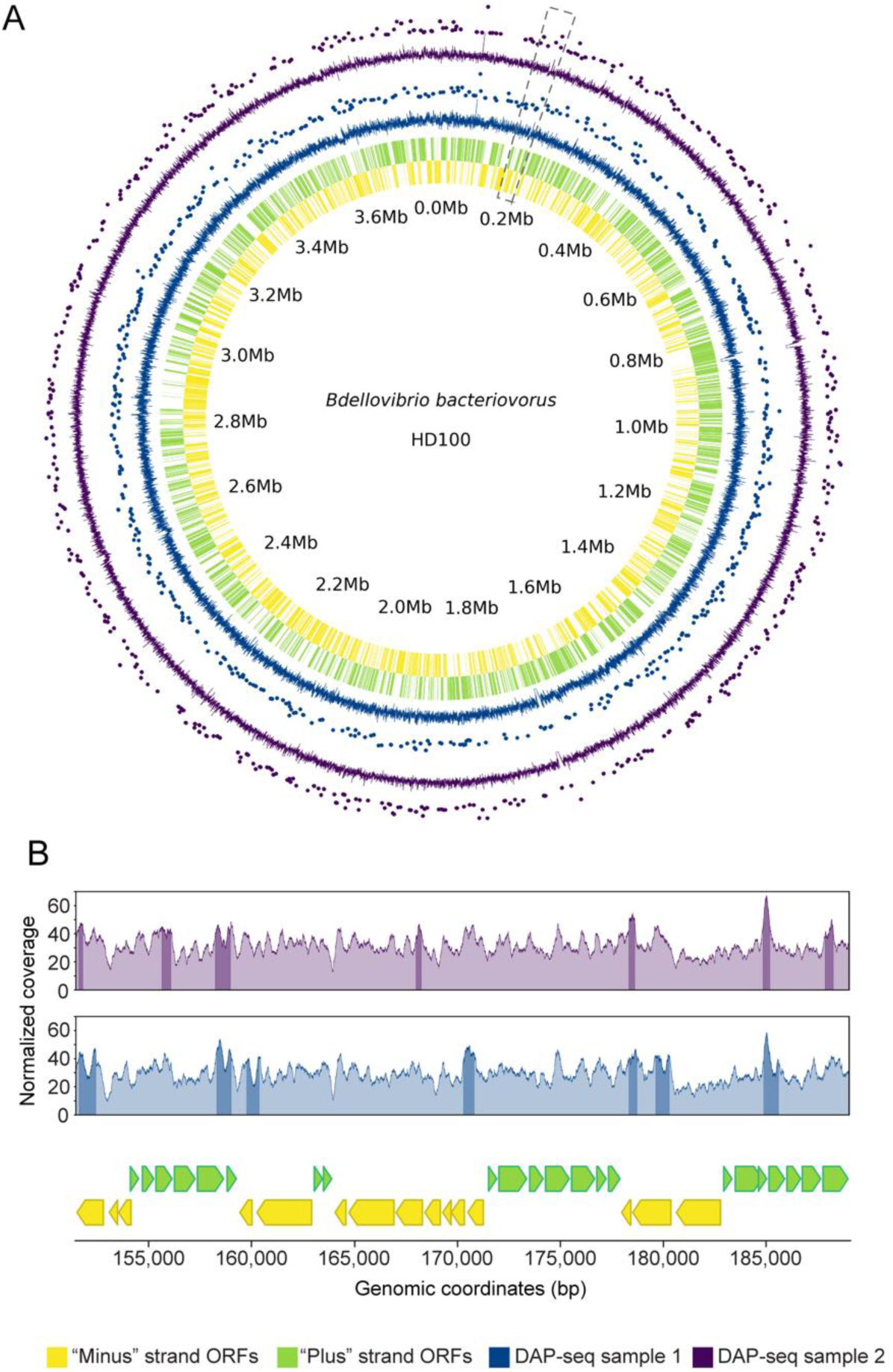
The HBb dimer binds DNA nonspecifically. **A.** DAP-seq data visualized using a circos plot of the *B. bacteriovorus* HD100 genome. The inner rug plots represent the “minus” (yellow) and “plus” (light green) strand ORFs. Each wedge represents the coordinates of a gene. Outer line plots represent the normalized read coverage values, and scatter plots above show the major peaks for the two DAP-seq samples (blue and purple). Peaks are shown as dots, where the strength of the peak signal correlates positively with the distance from the axis. The enlarged section shown in panel E is indicated by gray dashed line. **B.** Line plots showing the normalized read coverage values for a randomly selected section of the *B. bacteriovorus* HD100 genome encompassing the genomic region from 151,464 bp to 189,033 bp. The bottom plot shows the genomic context, and the top line plots depict the normalized read coverage values for each DAP-seq sample with the major peaks shaded darker. The color code corresponds to panel A.

## Discussion

Building on our recent discovery of bacterial histones (31), this study investigates the DNA binding and structuring properties of the histone protein HBb in *B. bacteriovorus*. We solved the crystal structure of HBb at a resolution of 1.06 Å, which notably, makes it the first structure of a histone protein at atomic resolution; the previous structure with the highest resolution (1.48 Å) was that of histone HMfA from *M. fervidus* (PDB: 1B67) (55). HBb features the characteristic histone fold and forms dimers. Most amino acid residues crucial for DNA binding in archaeal and eukaryotic histones are conserved in HBb. However, residues facilitating tetramerization in archaeal HMfB (17,43) are absent (Fig. 1A), suggesting that the dimeric state observed in both the crystal structure and biophysical data likely represent the physiological oligomer.

Employing a comprehensive array of assays, we demonstrate that HBb not only binds to DNA but also alters its topology. Our data indicate that HBb binds as individual dimers along the DNA and induces compaction through DNA bending, akin to the low-density, nonspecific binding mode of individual HU dimers (56). Importantly, MNase digestion assays and TPM measurements reveal that HBb does not compact DNA through wrapping, thereby ruling out nucleosome or hypernucleosome formation, a characteristic of canonical archaeal and eukaryotic histones (8,9,30). This is in agreement with a recent study that showed individual HBb dimers bind sequentially to DNA at increasing protein concentrations without forming hypernucleosomes (32). This binding mode is further supported by the HBb-DNA co-crystal structures presented in this study (32), yet a recently proposed model suggests that high-density binding of HBb would induce the formation of straight HBb-DNA filaments, contrasting our data that indicates compaction through bending. Such a straightening effect has been previously observed in the *E. coli* NAP HU at high density (57,58), but the physiological relevance of these structures remains unclear. It is possible that high-density binding of HU might occur on a local scale, driven by sequence determinants not yet identified. In the aforementioned study, an ensemble Förster Resonance Energy Transfer (FRET) assay was performed with a bi-terminally fluorescently labeled dsDNA molecule (32,59) and it confirmed that HBb does not trigger FRET, unlike the hypernucleosomal histone HTkA, which was used as a reference. This result supports the observation that HBb does not wrap DNA, though it offers only limited support for the notion that straight HBb-DNA filaments are formed. In fact, their MD simulations, instead, appear to suggest the binding and bending of DNA by individual HBb dimers (32).

Previous studies have reported that the expression of the *BD_RS00255* gene, encoding HBb, is upregulated during the replicative growth phase of *B. bacteriovorus*, a phase closely linked to prey consumption (60). This observation is corroborated by the most recent RNA-seq experiments, which also show that *BD_RS00255* is expressed exclusively during the replicative phase and not during the prey attack phase (Renske van Raaphorst and Géraldine Laloux, personal communication). Multiple independent attempts to disrupt the *BD_RS00255* gene have been unsuccessful, both by us and others (32), underscoring the essential role of HBb in the survival of *B. bacteriovorus*. DAP-seq analyses confirm that HBb binds to DNA across the entire genome, a pattern also observed in bacterial NAPs of the HU/IHF family. These proteins are encoded in 93% of bacterial genomes and are thought to play a role in controlling the dynamics and organization of the bacterial chromosome (61,62).

While our study primarily focuses on the bacterial histone HBb in *B. bacteriovorus*, the evolutionary implications of our findings merit further discussion. The presence of histone-like proteins in bacteria raises compelling questions about the evolutionary trajectory leading to the intricate histone structures and DNA-compacting capabilities observed in eukaryotes. One hypothesis is that the foundational aspects of histones and their DNA-binding abilities were already in place during the time of the Last Universal Common Ancestor (LUCA). These proteins may have initially served as ‘simple’ NAPs and subsequently underwent a series of evolutionary adaptations, enabling them to form nucleosomes and other higher-order chromatin structures. Such adaptations could have led to a gain of function, attributed to enhanced DNA affinity and (structural) binding specificity, priming the proteins for more efficient DNA packaging, organization, and specific regulatory roles. The presence of histones in a broad spectrum of deeply-branching bacterial lineages, combined with the conservation of key amino acid residues crucial for DNA binding across bacterial, archaeal, and eukaryotic histones, lends support to this notion. However, considering the infrequent occurrence of histones in bacteria and their sequence similarity to archaeal histones, we cannot rule out the possibility that these proteins were acquired by bacteria from archaea through horizontal gene transfer events. The future availability of additional bacterial histone sequences in molecular databases, as well as comparative genomic and functional studies, could illuminate the evolutionary pathways that have shaped histones across the domains of life.

In summary, our study establishes HBb as a unique bacterial histone with distinct DNA-structuring capabilities compared to eukaryotic, archaeal, and viral histones. Although its DNA-binding interfaces are similar to archaeal and eukaryotic histones, its architectural properties closely resemble those of HU/IHF family proteins. While HU/IHF family proteins also exist in *B. bacteriovorus*, they are present in significantly lower quantities compared to HBb (19). This supports the notion of functional redundancy among different bacterial NAPs, particularly in their global effects on chromosome organization. The indispensability of HBb for *B. bacteriovorus* survival suggests that bacterial histones, like their archaeal and eukaryotic counterparts, play critical roles in cellular functions.

## Data availability

Coordinates and structure factors of the crystal structures have been deposited in the PDB under accession codes 8CMP (free HBb), 9EZZ (HBb-DNA_1) and 9F0E (HBb-DNA_2).

The TPM, molecular dynamics data, and the supplementary videos are available from the 4TU repository (https://data.4tu.nl) with the DOI 10.4121/6c07e4bb-96f1-4ee0-88ef-6978bda27e5d. The DAP-seq raw data are deposited at Zenodo (https://zenodo.org/) and accessible with the DOI 10.5281/zenodo.10211306.

The analysis code for the molecular dynamics data is available under embargo from figshare (https://figshare.com) with the DOI 10.6084/m9.figshare.25574295.

## Acknowledgements

We are grateful to the staff of Beamline X10SA of the Swiss Light Source (PSI, Villigen, Switzerland) and beamline ID23-1 of the European Synchrotron Radiation Facility (Grenoble, France) for excellent technical support. We extend our thanks to Reinhard Albrecht and Antoine Schramm for assistance with crystallization, as well as crystallographic data collection and processing, and Kerstin Bär and Heidi Reichle for assistance with protein purification. We are grateful to Agnes Henschen, Min Zheng, Pengfei Liu (Dept. of Algal Development and Evolution, MPI for Biology, Tübingen, Germany) and Yihua Liu (Dept. of Microbiome Science, MPI for Biology, Tübingen, Germany) for their support with the DAP-seq methodology and data analysis. We thank the staff of the Genome Center, MPI for Biology Tübingen for coordination and help with next-generation DNA sequencing. We thank the ALICE HPC cluster at Leiden University for providing the necessary infrastructure to perform the molecular dynamics simulations.

