## Supplementary for "Bacterial histone HBb from *Bdellovibrio bacteriovorus* compacts DNA by bending"

**This file includes:**

SI Appendix, Figure S1 to S5

SI Appendix, Material and Methods

SI Appendix, Table S1 to S3

SI Appendix, References

### SI Appendix, Figures

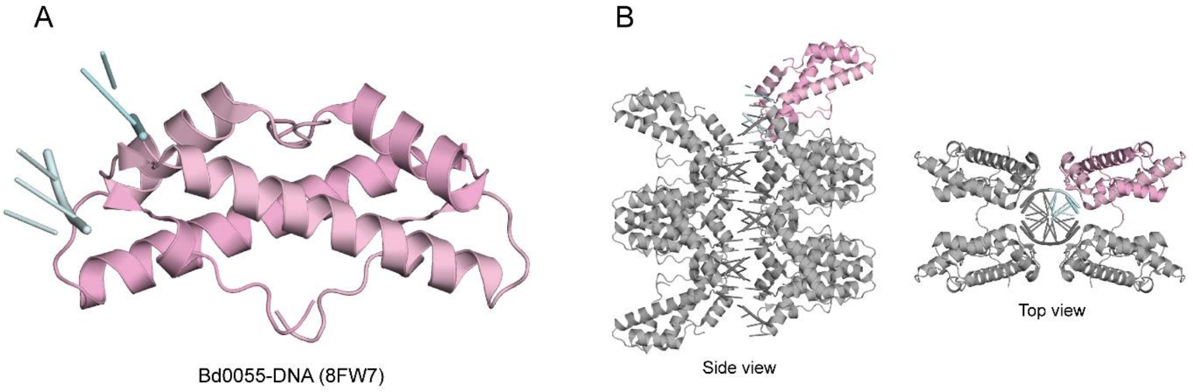

**Fig. S1. Crystal structure and crystal packing of DNA-bound HBb (Bd0055) structure (PDB: 8FW7) published by Hocher *et al.* (1). A.** The protein is shown in cartoon representation (in light pink) bound to DNA (in light cyan). **B.** The related crystal packing, visualized by selected symmetry mates (in gray) generated within 4 Å.

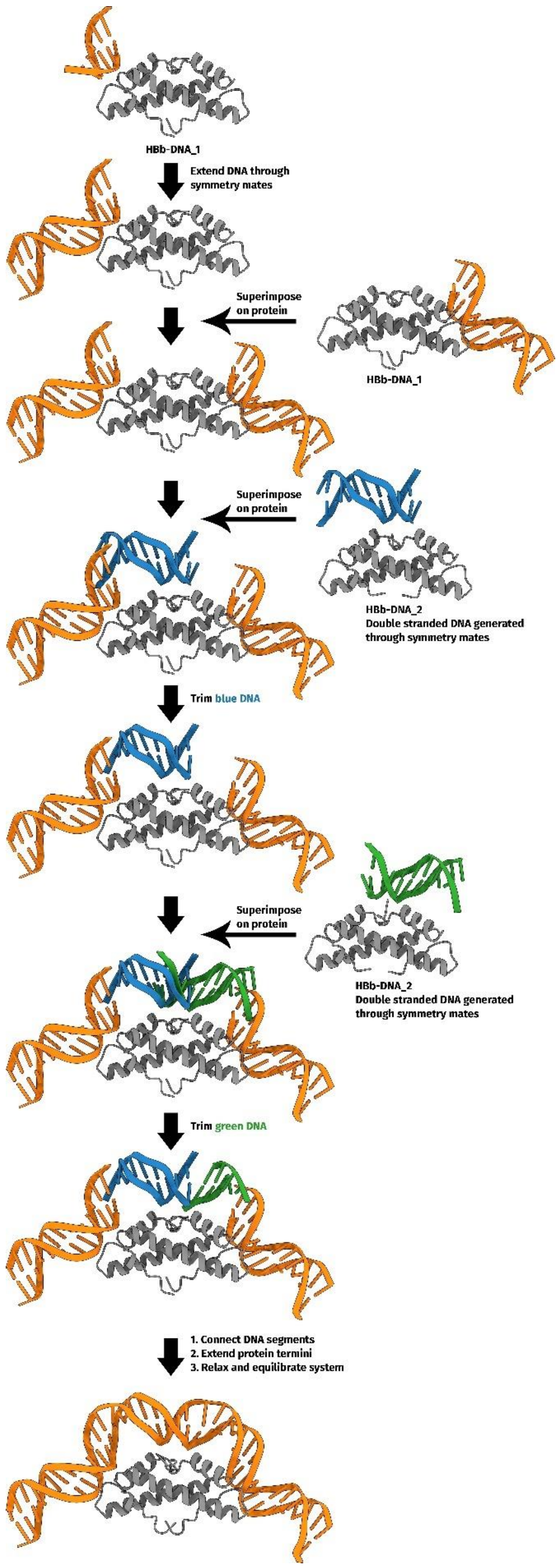

**Fig. S2. Method of generating a complete DNA molecule bent around HBb.**

The starting structure for the molecular dynamics simulations was constructed from the HBb-DNA_1 and 2 crystal structures. The DNA from HBb-DNA_1 is colored in orange. The DNA from HBb-DNA_2 is colored in blue and green. Protein is colored in gray.

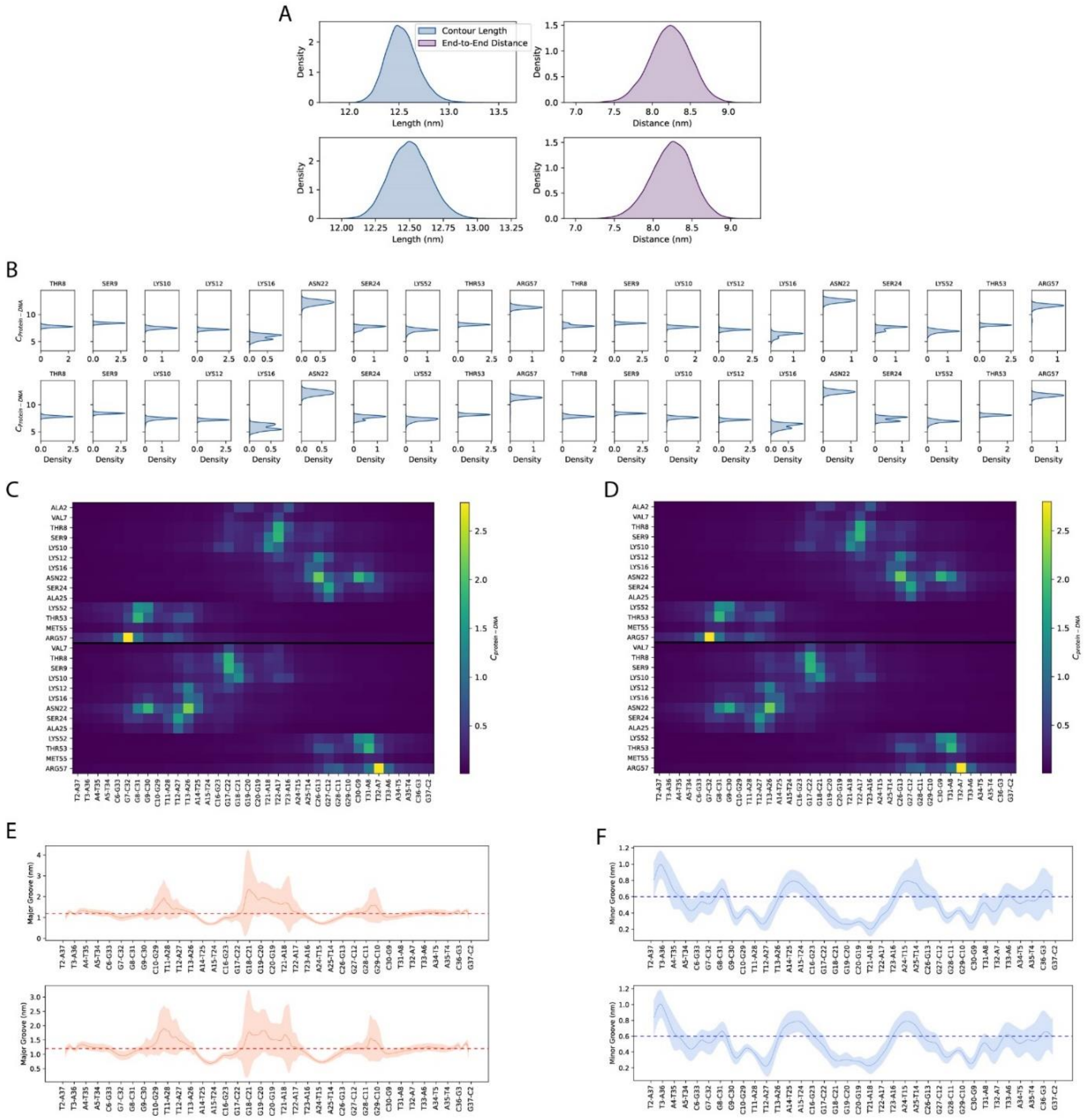

**Fig. S3. DNA structure and protein-DNA interaction analysis of the two 500 ns MD simulations. A.** Density estimates of the contour length and end-to-end distance for the DNA molecules of run 1 (top) and run 2 (bottom). **B.** Decomposed density estimates of the number of contacts of each protein residue with respect to the complete DNA of run 1 (top) and run 2 (bottom). Only residues with an average number of contacts above 5 are shown. The data for each monomer are concatenated. **C, D.** Heat map of the mean number of contacts of each protein residue with each base pair for run 1 (**C**) and run 2 (**D**). The data for each protein monomer is concatenated with a black dividing line separating the data of each monomer. **E, F.** Major (**E**) and minor (**F**) groove widths of the DNA molecules of run 1 (top) and run 2 (bottom). The shaded area represents one standard deviation. The normal groove width of B-DNA is indicated by the striped line.

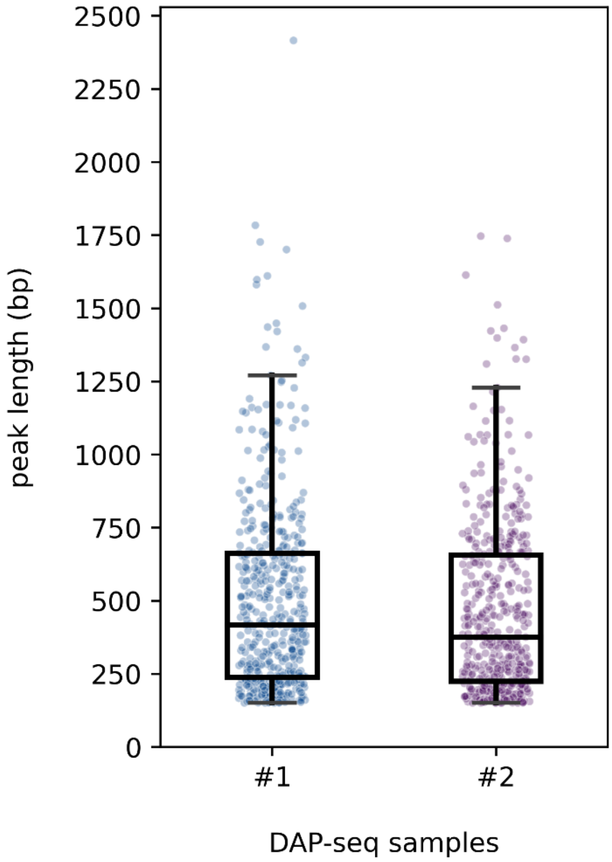

**Fig. S4.** Peak length distribution for DAP-seq samples 1 and 2.

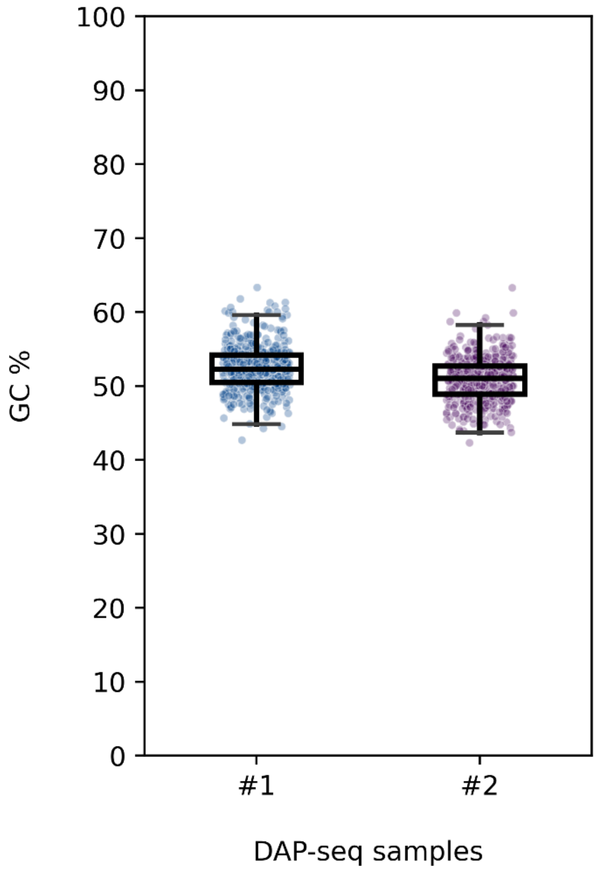

**Fig. S5.** GC content distribution of the genomic segments located between the coordinates of a given peak for DAP-seq samples 1 and 2.

### SI Appendix, Material and Methods

#### Bacterial strains and cultivation

Cloning procedures were performed in *Escherichia coli* Top10 cells. *E. coli* strain Mutant56(DE3) (2) was used as the expression host. *E. coli* cells were grown on Luria-Bertani (3) medium.

*B. bacteriovorus* HD100 (DSM 50701, DSMZ) was cultured at 30°C using *E. coli* Top10 as prey cells in PYE broth. Media was supplemented with antibiotics as required.

#### Cloning, plasmids, and synthetic DNA

The genes encoding HBb (*BD_RS00255,* old locus tag: *Bd0055*) and HMfB (GenBank accession number M34778.1) were codon-optimized and synthesized (Synbio Technologies; BioCat GmbH). The *hbb* gene was cloned into the expression vector pETHis1a (4) for recombinant expression of HBb (UniProt entry Q6MRM1) fused to an N-terminal $(histidine)_6$ tag and FLAG tag, respectively. The HMfB-encoding gene (UniProt entry P19267) was cloned in pET28a for expression with an N-terminal $(histidine)_6$ tag.

For chromosomal deletion of *BD_RS00255* in *B. bacteriovorus*, the suicide vector pT18mobSacB, kindly gifted by Brian Kvitko (Addgene plasmid # 72648; http://n2t.net/addgene:72648; RRID: Addgene 72648), was used. The kanamycin resistance ($Km^r$) cassette was amplified from plasmid pT2SK, provided as a gift from Shelley Copley (Addgene plasmid # 59383; http://n2t.net/addgene:59383; RRID:Addgene_59383). Vector pUC18 was used for subcloning. Synthetic oligonucleotides (Merck) used in this study are listed in Table S1.

#### Deletion of *BD_RS00255*

For deletion of *BD_RS00255* in *B. bacteriovorus* HD100, the genomic regions upstream and downstream of *BD_RS00255* (nucleotides 47964-49345 and 49567-50958 of the complete genome of *B. bacteriovorus* strain HD100; segment 1/11, Sequence ID: BX842646.1) were amplified by PCR using specific primers (Table S1) and cloned into pUC18. The $Km^r$ cassette was amplified by PCR from vector pT2SK and inserted into pUC18 between both fragments. A fragment comprising the $Km^r$ cassette flanked by the *BD_RS00255* upstream and downstream genomic region was amplified by PCR and subcloned into the suicide vector pT18mobSacB. The resulting plasmid was introduced into *B. bacteriovorus* via conjugation transformation. Chromosomal integration of the plasmid by homologous recombination was verified by PCR and sequencing. To select for a second crossover event resulting in the replacement of *BD_RS00255* by the $Km^r$ cassette, cells were grown on medium containing 5-10% sucrose.

**Protein expression and purification**

For expression of FLAG-HBb, His-HBb, and HMfB in *E. coli* Mutant56(DE3), cells that were transformed with the respective plasmids encoding were cultivated in LB broth, supplemented with kanamycin at 37°C. At an $OD_{600}$ of 0.5, IPTG was added at a final concentration of 1 mM to induce protein expression. Following further cultivation at 25°C for 16 h for FLAG-HBb, and at 37°C for 4 hours for His-HBb and HMfB, respectively, cells were harvested by centrifugation. All purification steps were performed at 4°C. For HBb proteins, all buffers were supplemented with 2-mercaptoethanol (2.5 mM) to prevent cysteine disulfide bridge formation. Cell pellets were resuspended in a buffer containing 20 mM Tris, pH 8.0, 300 mM NaCl and 10 mM imidazole, supplemented with protease inhibitor mix (cOmplete™, EDTA-free Protease Inhibitor Cocktail, Roche), 0.1 mM PMSF, 3 mM $MgCl_2$ and DNase. Cells were lysed by sonication for 15 min. Cell debris and membranes were pelleted at 95,000 x g for 45 min. The supernatant was filtered with a 0.45 µm filter and applied on a 5 mL HisTrap column (Cytiva). The bound proteins were eluted with a linear gradient of 10-500 mM imidazole in the aforementioned buffer. For His-HBb, the histidine tag was cleaved with TEV protease. Following a second HisTrap column to remove the histidine-tagged TEV, HBb was purified to homogeneity by gel filtration chromatography on a Superdex 75 column (Cytiva) equilibrated with SEC buffer (20 mM Tris, pH 8.0, 150 mM NaCl). Protein purity was assessed by SDS-PAGE (15%) and BN-PAGE (4-16%, ThermoFisher Scientific). The protein concentration was determined using the BCA protein assay (ThermoFisher Scientific). Purified HBb was dialyzed against 50 mM Tris, pH 7.0, 75 mM KCl and 10% Glycerol for further assays.

**Circular dichroism (CD) spectroscopy**

HBb sample was dialyzed in a dialysis buffer (10 mM Tris, pH 8.0, 100 mM KF and 1 mM 2-mercaptoethanol) overnight, and then diluted with the dialysis buffer to a final concentration of 25 µM. CD spectra were measured with a Jasco J-810 spectrometer (JASCO) in a wavelength range of 190-240 nm, using a cuvette with a path length of 1 mm and a read speed of 100 nm/min. A total of ten single spectra were recorded and averaged. Thermal stability of HBb was analyzed measuring the ellipticity (θ) at 220 nm over a temperature gradient of 20-100°C applying a ramp of 1°C/min, a data pitch of 0.5, and a response time of 1 s. Data analysis, including blank subtraction and curve smoothing, was performed using the Spectra Manager software suite (JASCO). CD spectrum and the melting curve were plotted using Prism 9 (GraphPad).

**Size exclusion chromatography coupled with multi-angle light scattering (SEC-MALS)**

SEC-MALS of HBb was performed in SEC buffer. HBb at a concentration of 2.7 mg/mL was incubated on ice for 20 min before loading onto a Superdex 75 Increase 10/300 GL column

(Cytiva), equilibrated with the corresponding buffer. The run was performed at a flow rate of 0.5 mL/min on a 1260 Infinity II HPLC system (Agilent) coupled to a miniDAWN TREOS and Optilab T-rEX refractive index detector (Wyatt Technology). HBb was detected at 215 nm and measured in triplicate. Molar mass distributions were calculated using the ASTRA v.7.3.0.18 software suite (Wyatt Technology).

**Crystallization, data collection, and structure determination**

Crystallization trials were performed by mixing 300 nL of protein with 300 nL of reservoir solution in 96-well sitting-drop vapor-diffusion plates using commercially available screens with reservoir volumes of 100 µL. In free form, HBb was set up at a concentration of 2.7 mg/ml in 20 mM Tris, pH 8.0, 150 mM NaCl, 1 M Urea, and best crystals were obtained with a reservoir solution containing 0.1 M $CH_3COONa$, pH 4.5 and 25% (w/v) PEG 3350. For co-crystallization, the protein was incubated with the randomized double-stranded DNA fragment 20 bp-GC50 (Table S1) at a volume ratio of 1:1 at 37 °C for 10 min, and aggregates removed by centrifugation after cooling to room temperature. Best crystals were obtained with a protein solution containing 800 µM HBb and 200 µM DNA and reservoir containing 0.1 M HEPES, pH 7.0, 0.2 M LiCl and 20% (w/v) PEG 6k (HBb-DNA_1) and protein solution containing 600 µM HBb and 150 µM DNA and a reservoir of 0.2 M $(NH_4)_2SO_4$, 0.1 M $CH_3COONa$, 22% (w/v) PEG 4k (HBb-DNA_2). For cryo protection, crystals were transferred to droplets of their reservoir solution spiked with 10% (v/v) PEG 400 (free HBb), 20% (v/v) PEG 400 (HBb-DNA_1) or 15% (v/v) PEG 200 (HBb-DNA_2), loop-mounted, and flash-cooled in liquid nitrogen. Data were collected at beamline X10SA of the Swiss Light Source (Villigen, Switzerland) (free HBb) and beamline ID23-1 of the European Synchrotron Radiation Facility (Grenoble, France) (HBb-DNA_1, HBb-DNA_2) at 100 K, using EIGER X 16M hybrid pixel detectors (Dectris Ltd., Switzerland). Data were reduced, processed and scaled using XDS (5).

The structure of free HBb was solved by molecular replacement (MR) using Phaser (6) and HMk (PDB: 1F1E) as a search model, locating one monomer in the asymmetric unit (AU). The structure was rebuilt and refined in cycles with manual modelling in Coot (7) and refinement with REFMAC5 (8). DNA-bound structures were solved by MR using MOLREP (9) and the refined free HBb coordinates as a model, locating one dimer in the AU for both structures. After initial rigid body refinement with REFMAC5 (8), electron density for double-stranded DNA fragments became apparent in both structures in different locations, which were modeled manually based on the DNA sequence. All structures were finalized in cycles of manual modelling in Coot (7) and refinement in REFMAC5 (8). Data processing and refinement statistics are listed in Table S3, and the coordinates and structure factors deposited in the PDB under accession codes 8CMP (free HBb), 9EZZ (HBb-DNA_1) and 9F0E (HBb-DNA_2).

### Microscale thermophoresis (MST)

To determine the DNA binding affinity of HBb, Cy5-labelled 80 bp DNA (10,11) and 80 bp-GC50 oligonucleotides were annealed as described. A dilution series of label-free HBb was prepared in 20 mM Tris, pH 8.0, 150 mM NaCl and 2.5 mM 2-mercaptoethanol and titrated against the annealed DNA fragments at a concentration of 10 nM. The HBb-DNA mixtures were incubated at 37°C for 10 min, centrifuged to remove precipitates, and loaded into Monolith NT premium capillaries (MO-K025, NanoTemper Technologies) for measurement. HMfB was used as a control. Similarly, a dilution series of the label-free HMfB was titrated against both DNA fragments at the concentration of 20 nM. HMfB-DNA mixtures were incubated at RT for 5 min, centrifuged and loaded into Monolith NT capillaries (MO-K022). All measurements were repeated in triplicate and performed at 25°C using the Monolith NT.115 instrument with a Nano RED Detector and MST power set to medium. MST data were analyzed by fitting them to a $K_d$ model using MO.Control (NanoTemper Technologies).

### DNA binding analysis

*Electrophoretic mobility shift assay (EMSA)*

For *in vitro* DNA-binding tests, an 80 bp DNA fragment (10,11) was used. For annealing, complementary oligonucleotides were incubated at 95°C for 5 min, followed by slow cooling to room temperature. The annealed 80 bp DNA was mixed with protein at different ratios in 25 mM Tris, pH 8.0, 50 mM NaCl, 50 mM KCl and incubated at 37°C for 10 min. Samples were separated on a 6% DNA retardation gel (ThermoFisher Scientific) at 100 V and DNA was visualized by SYBR Gold (ThermoFisher Scientific) staining.

To approximate the length of DNA fragments required for protein binding, GeneRuler Ultra Low Range DNA Ladder (ThermoFisher Scientific) was mixed with HBb or HMfB in 50 mM Tris, pH 7.0, 75 mM KCl at indicated ratios and incubated at room temperature for 30 min. Glycerol was added to a final concentration of 10%, and samples were separated on a self-made 10% polyacrylamide gel in TAE buffer at 120 V. DNA was stained with GelRed (Biotium) and visualized using Gel Doc XR+ imaging system (Bio-Rad Laboratories).

*Micrococcal nuclease (MNase) digestion assay*

For HBb-DNA complex formation, 1440 ng HBb was mixed with 900 ng 685 bp DNA in 50 mM Tris, pH 7.0, 75 mM KCl, and incubated at room temperature for 30 min. Then, indicated amounts of MNase (New England BioLabs) were added and incubation was continued in digestion buffer, provided by the manufacturer, for 12 min at 37°C in a total reaction volume of 80 µL. The reaction was stopped with EDTA and SDS at a final concentration of 95 mM and 0.5%, respectively. The protected DNA was recovered by phenol/chloroform extraction, followed by ethanol precipitation. The purified DNA was dissolved in 5 µL nuclease-free $H_2O$

and separated on a self-made 10% polyacrylamide gel in TAE buffer at 120 V for 1 h. To visualize the DNA, the gel was stained with GelRed (Biotium) and imaged using Gel Doc XR+ imaging system (Bio-Rad Laboratories). For the positive control HMfB, 540 ng protein was incubated with 900 ng 685 bp DNA and the assay was performed as described.

*DNA topology assay*

*E. coli* DH5α cells were freshly transformed with plasmid pUC19 and grown to an $OD_{600}$ of 2 to isolate the plasmid using GeneJET Plasmid Miniprep Kit (ThermoFisher Scientific). Topologically relaxed pUC19 was obtained by nicking with endonuclease Nb.BsrDI (New England BioLabs) followed by covalent closure with T4 DNA ligase (ThermoFisher Scientific). Successful relaxation was confirmed by agarose gel electrophoresis. In the assay, 200 ng of relaxed pUC19 was mixed with 100 ng, 200 ng, and 400 ng HBb and HMfB, respectively, in 50 mM Tris, pH 7.0, 75 mM KCl. After 30 min incubation at room temperature, 7.5 U of topoisomerase I (type IB) (ThermoFisher Scientific) was added and incubation continued for 60 min at 37°C in a total reaction volume of 50 µL. Pure DNA was obtained by phenol/chloroform extraction followed by ethanol precipitation. Samples were separated on a 1% TAE agarose gel. To visualize different topological states of the plasmid, DNA was stained with GelRed (Biotium) and imaged using Gel Doc XR+ imaging system (Bio-Rad Laboratories).

*Ligase-mediated circularization assay*

The ligase-mediated circularization assay was performed with a linear DNA fragment of 240 bp comprising the first 240 bp of the DNA substrate used in the TPM experiments. 380 ng DNA was mixed with 108 ng, 216 ng and 432 ng HMfB, and 288 ng, 576 ng and 1152 ng HBb in 50 mM Tris, pH 7.0, 75 mM KCl. Following incubation at RT for 30 min, $MgCl_2$, ATP, DTT, and T4 DNA ligase were added to final concentrations of 10 mM, 1 mM, 10 mM, and 0.2 U/µL, respectively, in a total volume of 100 µL and incubated for 24 hours at room temperature. The DNA was purified by phenol/chloroform extraction and ethanol precipitation. One third of purified DNA was subjected to T5 exonuclease (New England BioLabs) treatment in CutSmart™ buffer (New England BioLabs) for 60 min at 37°C. DNA samples were separated on a 2% TAE agarose gel; stained with GelRed (Biotium) and visualized using Gel Doc XR+ imaging system (Bio-Rad Laboratories).

*Tethered Particle Motion (TPM) experiments*

TPM experiments were performed as previously described (12) using 50 mM Tris, pH 7.0, 75 mM KCl as buffer. A standard deviation cutoff of 8% and an anisotropic ratio cutoff of 1.3 were used to select single-tethered beads. Measurements at each HBb concentration were done in triplicate. Means and standard deviations of the individual measurement series for each HBb concentration were calculated by maximum likelihood estimation assuming a normal distribution. Outliers with a Z-score >3 or <-3 were not considered for fitting. The “line to guide the eye” was generated by fitting the means to a logistic function. A custom Python script was used for fitting and plotting the TPM data. For plotting, the means of the three individual measurements were averaged for each measured concentration and the standard deviations were error-propagated ($Std(\bar{X}) = \sqrt[2]{\frac{\sum_{i=1}^{n} Var(X_i)}{n^2}}$).

### DAP-seq

*Library construction and DNA precipitation*

Genomic DNA (gDNA) libraries of *B. bacteriovorus* were constructed with an average fragment length of 200 bp as described (13). For DNA precipitation, 5 µg of purified FLAG-HBb was diluted in Binding buffer (PBS supplemented with 1 mM 2-mercaptoethanol) to a final volume of 400 µL and incubated with 25 µL of pre-washed Anti-FLAG M2 magnetic beads (Sigma-Aldrich) for 60 min at room temperature in an orbital shaker. Subsequently, the beads were washed 5 times with Binding buffer supplemented with 0.005% NP-40, followed by three wash steps with non-supplemented Binding buffer. Washed beads were resuspended in 80 µL of Binding buffer containing 1 µg of adaptor-ligated gDNA library and incubated for 60 min at room temperature in an orbital shaker. The beads were washed a total of eight times with 200 µL of Binding buffer, with the first 5 washing steps conducted in the presence of 0.005% NP-40. Bound DNA-protein complexes were eluted from the beads by incubation with 30 µL 3xFLAG peptide (100 ng/µL, Sigma-Aldrich) at room temperature for 30 min in an orbital shaker. Eluted DNA was amplified using different pairs of indexed primers for each sample. Removal of primer dimers and size selection of PCR products was performed using AMPure XP beads (Beckman Coulter) according to the instruction and subsequent elution in 20 µL 0.1x TE buffer. The concentration of each library was measured using Qubit dsDNA HS assay kit (Invitrogen) and the average size of each library was determined by Bioanalyzer DNA HS kit (Agilent). Libraries were diluted to a concentration of 2 nM and pooled for sequencing. The negative control experiment was conducted by incubating the gDNA library with beads in the absence of protein. All DAP-seq experiments were repeated twice.

*Next-generation Sequencing*

Sequencing was performed using an Illumina NextSeq 2000 instrument. An average of 2 million pair-end 151-bp reads per sample were generated.

*Data analysis*

The paired-end FASTQ files were processed with the nf-core/chipseq (version 2.0.0) pipeline (14), under Nextflow (version 23.04.2). The *B. bacteriovorus* HD100 genome (assembly ASM19617v1, RefSeq accession GCF_000196175.1) was used as reference, and the respective nucleotide FASTA, and GTF annotation files were provided as inputs to the pipeline. The chosen configuration profile was that of Singularity, and the effective genome size parameter was set to 3,782,950. All other parameters were left to default. The BAM files generated by the nf-core/chipseq pipeline (".mLb.clN.sorted.bam" suffix) were provided as input to MACS (version 3.0.0b3) (15). MACS was run without building a model, with a shift-size of 151 bp, the "broad" flag set, and the effective genome size parameter set to 3,782,950. All other parameters were left to default. The BED files generated by MACS ("peaks.gappedPeak" suffix), were analyzed with pandas (version 1.5.3) (16). The GC content of the genomic segments located between the coordinates of a given peak was calculated with BioPython (version 1.81) (17), using these BED files, and the nucleotide FASTA for the genome described above as inputs. The bigWig files generated by the nf-core/chipseq pipeline were analyzed with Bioframe (version 0.4.1) (18), in conjunction with the genomic features table file, and GenBank flatfile pertaining to the genome described above. Normalized read coverage values, peak coordinates, peak signal values, peak length, GC content, and genomic coordinates were visualized with pyCircos (github.com/ponnhide/pyCircos) and Matplotlib (version 3.7.1) (19). All Python packages were installed and used under Python 3.10.12.

**Molecular dynamics (MD) simulations**

For MD simulations, we used GROMACS (20) with the AMBER ff14sb-ParmBSC1 force field (21-23). All systems were solvated in a dodecahedron box with a distance of at least 1.0 nm to the box boundary and filled with SPC/E water molecules (24). Water molecules were replaced at random with $Na^+$ and $Cl^-$ ions to charge-neutralize the system. 75 mM of $Na^+$ and $Cl^-$ ions were additionally added. Energy minimization was performed using the steepest descent method for 5000,000 steps until the largest force was below 1000.0 kJ/mol/nm. To equilibrate the solvent and ions, heavy atoms were position constrained for 100 ps at 310 K and 1 bar. The cutoff for van der Waals interactions was 1.1 nm. Electrostatic interactions beyond a cutoff of 1.1 nm were treated with the particle-mesh Ewald method using a grid spacing of 0.16 nm. Temperature and pressure were kept constant with the V-rescale thermostat (Bussi-Donadio-Parrinello thermostat) (25) and the Parrinello-Rahman barostat (26), respectively. Bonds were constrained with LINCS (27) and simulations were conducted in time steps of 2 fs.

The starting structure for the MD, with DNA bent around the HBb dimer, was constructed from the HBb-DNA_1 and HBb-DNA_2 crystal structures (Fig. S2). The DNA at the l1-l2 loops of the dimer comes from the HBb-DNA_1 crystal structure. The DNA from HBb-DNA_1 was extended through crystal symmetry mates. The DNA between the two l1-l2 loops was filled in with the DNA from the HBb-DNA_2 crystal structure by superimposition on the protein, followed by trimming of the DNA. All DNA segments were connected together and the sequence was changed to the sequence of the 20 bp-GC50 fragment used for crystallization. For the final structure, the protein structure of HBb-DNA_2 was used and the missing terminal protein residues were filled in based on an AlphaFold prediction of the HBb dimer, generated with ColabFold (28,29). All protein superimpositions, trimming of the DNA, and the changes to the DNA sequence were conducted in ChimeraX v1.7.1 (30). Crystal symmetry mates were generated in Open-Source PyMOL v2.5.0 (31,32).

Snapshots were collected every 40 ps from the last 100 ns of both runs and clustered on RMSD of all protein and DNA atoms with a cutoff value of 0.2 Å in GROMACS with the gromos algorithm (33). Clustering resulted in 60 clusters, the largest of which contained 1873 snapshots (37% of all snapshots analysed during clustering). From the most abundant cluster, the central structure, i.e., the structure with the smallest average RMSD among all other structures of the cluster, was used as the representative structure of the MD in Figure 5C and 5D.

The atoms involved in protein-DNA contacts were identified as described in a previous study by van Heesch et al. (34).

The bending angle of the DNA was computed by first converting the DNA base pairs into a rigid body model (35), which defines mean reference frames for each base pair. Then, after defining vectors through the origins of the mean reference frames of the second base pair and center base pair, and the second to last base pair and center base pair, the bending angle was computed as 180 ° minus the inverse cosine of the dot product of these vectors. For the bending angle of HMfB, the same procedure was performed on the crystal structure of HMfB bound to DNA (PDB: 5T5K) (11).

Data was plotted in Python with the Matplotlib package (19,32).

### SI Appendix, Tables

#### Table S1

#### Oligonucleotides used in this work

| Oligo name | Sequence (5’-3’) | Application |
|---|---|---|
| **hbb up fwd (SalI)** | 5’-CTAGTCGACCATGTGTGAAGACCCC ATCAATC-3’ | *hbb* deletion, amplification of *BD_RS00255* upstream region |
| **hbb up rev (Acc65I)** | 5’-CTAGGTACCCCATGTTAATCGAACAG ATAAAAG-3’ | *hbb* deletion, amplification of *BD_RS00255* upstream region |
| **hbb dwn fwd (Acc65I)** | 5’-CTAGGTACCGAAAATTCGTTCAGCAT TGGTTC-3’ | *hbb* deletion, amplification of *BD_RS00255* downstream region |
| **hbb dwn rev (SacI)** | 5’-CTAGAGCTCCTTGGGTGTGAAGATCA TCTCTTG-3’ | *hbb* deletion, amplification of *BD_RS00255* downstream region |
| **Km fwd (Acc65I)** | 5’-CTAGGTACCCTCTGATGTTACATTGC ACAAGATAAAA-3’ | *hbb* deletion, amplification of $Km^r$ cassette |
| **Km rev (Acc65II)** | 5’-CTAGGTACCTCCTGCGTTACGCCCC GCCCTGC-3’ | *hbb* deletion, amplification of $Km^r$ cassette |
| **pUC/M13 fwd (XbaI)** | 5’-GCTATCTAGAGTAAAACGACGGCCA GTGCC-3’ | *hbb* deletion, amplification of $Km^r$ cassette flanked by *BD_RS00255* up- and downstream regions |
| **pUC/M13 rev (XbaI)** | 5’-GCTATCTAGACAGGAAACAGCTATGA CCATG-3’ | of $Km^r$ cassette flanked by *BD_RS00255* up- and downstream regions |
| **20 bp-GC50-F** | 5’-TTAAAGCCCGTTAAAGCCCG-3’ | Co-crystallization of HBb and DNA |
| **20 bp-GC50-R** | 5’-CGGGCTTTAACGGGCTTTAA-3’ | Co-crystallization of HBb and DNA |
| **80 bp-DNA-Fwd [Cy5]** | 5’-[cyanine5]CCGTACTGTCGTCTGCGGC CTTTGATTATCAATTAAAGCGTTCTACG GCGTTTTTGATCGCTCAACGTGCGGAG CTAGAT-3’ | MST |
| **80 bp-DNA-Fwd** | 5’-CCGTACTGTCGTCTGCGGCCTTTGAT TATCAATTAAAGCGTTCTACGGCGTTTT TGATCGCTCAACGTGCGGAGCTAGAT-3 ’ | EMSA |
| **80 bp-DNA-Rev** | 5’-ATCTAGCTCCGCACGTTGAGCGATCA AAAACGCCGTAGAACGCTTTAATTGATA ATCAAAGGCCGCAGACGACAGTACGG-3’ | EMSA, MST |

| | | |
|---|---|---|
| **80 bp-GC50-Fwd [Cy5]** | 5’-[cyanine5]TTAAAGCCCGTTAAAGCCCGTTAAAGCCCGTTAAAGCCCGTTAAAGCCCGTTAAAGCCCGTTAAAGCCCGTTAAAGCCCG-3’ | MST |
| **80 bp-GC50-Rev** | 5’-CGGGCTTTAACGGGCTTTAACGGGCTTTAACGGGCTTTAACGGGCTTTAACGGGCTTTAACGGGCTTTAACGGGCTTTAA-3 | MST |
| **240 bp-DNA-Fwd** | 5’-[Phos]TTACTTTCACCAGCGTTTCTGGGTGAGCAAAAACAG-3’ | Ligase-mediated circularization assay |
| **240 bp-DNA-Rev** | 5’-[Phos]TGGTTTCTTAGACGTCAGGTGGCACTTTTCGG-3’ | Ligase-mediated circularization assay |
| **685 bp-DNA-Fwd [Biotin]** | 5’-[Biotin]TTACTTTCACCAGCGTTTCTGGGTGAGCAAAAACAG-3’ | TPM |
| **685 bp-DNA-Rev [DIG]** | 5’-[DIG]CCAAGTAGCGAAGCGAGCAGGACTGGGCGG-3’ | TPM |
| **685 bp-DNA-Fwd** | 5’-TTACTTTCACCAGCGTTTCTGGGTGAGCAAAAACAG-3’ | MNase assay |
| **685 bp-DNA-Rev** | 5’-CCAAGTAGCGAAGCGAGCAGGACTGGGCGG-3’ | MNase assay |

**Table S2**

**Binding affinities ($K_d$) of HBb and HMfB to DNA substrates.**

| Protein | DNA substrate | $K_d$ ± SD (μM) |
|---|---|---|
| HBb | 80 bp DNA | 4.84 ± 0.97 |
| | 80 bp-GC50 DNA | 11.1 ± 1.07 |
| HMfB | 80 bp DNA | 0.83 ± 0.27 |
| | 80 bp-GC50 DNA | 3.92 ± 1.16 |

**Table S3**

**Data collection and refinement statistics of free HBb, HBb-DNA_1 and HBb-DNA_2**

Values for the outer shell are given in parentheses.

| | HBb | HBb-DNA_1 | HBb-DNA_2 |
|---|---|---|---|
| **Data collection** | | | |
| Space group | $P2_12_12$ | $P2_12_12_1$ | C2 |
| Cell dimensions | | | |
| *a*, *b*, *c* (Å) | 34.47, 56.35, 26.70 | 32.32, 34.22, 151.83 | 108.70, 34.76, 56.43 |
| α, β, γ (°) | 90, 90, 90 | 90, 90, 90 | 90, 100, 90 |
| Resolution range (Å) | 28.17–1.06 (1.13–1.06) | 31.63-1.95 (2.07-1.95) | 42.49-1.85 (1.96-1.85) |
| Completeness (%) | 96.3 (79.4) | 99.9 (99.8) | 93.0 (95.2) |
| Redundancy | 10.8 (3.88) | 7.89 (8.33) | 4.68 (4.70) |
| $\langle I/\sigma(I)\rangle$ | 23.80 (1.66) | 11.3 (0.86) | 9.59 (2.18) |
| $R_{meas}$ | 0.047 (0.921) | 0.104 (2.16) | 0.110 (0.637) |
| **Refinement** | | | |
| No. of reflections, working set | 20904 | 11629 | 15089 |
| No. of reflections, test set | 1161 | 645 | 843 |
| Final $R_{cryst}$ | 0.142 | 0.221 | 0.249 |
| Final $R_{free}$ | 0.169 | 0.268 | 0.267 |
| R.m.s. deviations | | | |
| Bonds (Å) | 0.019 | 0.005 | 0.008 |
| Angles (°) | 1.421 | 1.271 | 1.297 |
